# Evolutionary Dynamics of Non-Coding Regions in Pancreatic Ductal Adenocarcinoma

**DOI:** 10.1101/2020.09.11.294389

**Authors:** Akimasa Hayashi, Yu-jui Ho, Alvin P. Makohon-Moore, Amanda Zucker, Jungeui Hong, Johannes G. Reiter, Jinlong Huang, Lance Zhang, Marc A. Attiyeh, Priscilla Baez, Rajya Kappagantula, Jerry P. Melchor, Eileen M. O’Reilly, Nicholas D. Socci, Shinya Oki, Scott W. Lowe, Christine A. Iacobuzio-Donahue

## Abstract

While the non-coding genome appears to play a role, the dynamic nature of noncoding alterations with respect to clonal progression of solid tumors remains unexplored. To address this gap in knowledge we performed multiregional whole genome sequencing and clonal analysis to elucidate the evolutionary dynamics of non-coding regions in pancreatic cancer relative to those of the coding genome. We find that the mutational burden of noncoding DNA is higher than coding DNA. However, when noncoding DNA was segregated into enhancer and non-enhancer regions, enhancers were more similar to coding DNA. Mutational signatures of noncoding and coding DNA further revealed the similar mutational spectra of enhancers to coding DNA whereas the mutational spectra of non-enhancer, noncoding DNA had an entirely different pattern. These findings shed light on the role of noncoding DNA in pancreatic cancer.

## Introduction

Despite the wealth of data pertaining to the biology and genetics of pancreatic ductal adenocarcinoma (PDAC), this solid tumor remains one of the most lethal tumor types (Kamisawa *et al*., 2016)(Siegel, Miller and Jemal, 2019). Large scale whole exome sequencing studies have revealed the recurrent genomic features of PDAC that target a defined number of core pathways (Jones *et al*., 2008)(Bailey *et al*., 2016)(Biankin *et al*., 2012)(N Waddell *et al*., 2015)(Witkiewicz *et al*., 2015). High frequency driver gene mutations in PDAC include *KRAS, TP53, CDKN2A, SMAD4* which play a major role in clonal expansions during pancreatic carcinogenesis and influence metastatic propensity. Subsequent subclonal genetic events are also key for tumor progression and metastasis, for example copy number gains of *cMYC* (Hayashi *et al*., 2020). Copy number alterations during clonal progression have been reported for multiple tumor types to date (Jamal-Hanjani *et al*., 2017)(Gerlinger *et al*., 2012).

Emerging data indicate the noncoding genome also plays a role in cancer. For example, increased long interspersed element 1 (LINE-1 or L1) activity has been was associated with multiple tumor types where it may contribute to somatic structural variations in known driver genes (Rodriguez-Martin *et al*., 2020). Mutations in promoter regions are also known as a functional target of mutational events. *TERT* promoter is one of the most notable non-coding functional mutations, first described in malignant melanoma (Huang *et al*., 2013) (Horn *et al*., 2013) and confirmed by comprehensive re-analyses of 863 individual whole genome sequence data across over 25 types of cancer (Weinhold *et al*., 2014). In PDAC, Feigin et al found an impact of mutations in promoter regions, specifically cis-regulatory regions of genes associated with transcriptional regulation (Feigin *et al*., 2017). Fewer studies have focused specifically on enhancer mutations (Weinhold *et al*., 2014)(Perera *et al*., 2016). Recently the ICGC/TCGA Pan-Cancer Analysis of Whole Genomes Consortium suggested that mutations in enhancer regions may play an important role in transcriptional changes through transcription factor dynamics (Campbell *et al*., 2020) (Andersson and Sandelin, 2020). Furthermore, in some tumor types such as bladder (Wu *et al*., 2019) and breast cancer (Rheinbay *et al*., 2017) enhancer mutations are thought to be driver mutations, a, observation also supported by pan-cancer analyses (Fredriksson *et al*., 2014)(Shuai *et al*., 2020).

While the non-coding genome appears to play a role, the dynamic nature of noncoding alterations with respect to clonal progression of solid tumors remains unexplored (Mimori *et al*., 2018). To address this gap in knowledge we performed multiregional whole genome sequencing and clonal analysis to elucidate the evolutionary dynamics of non-coding regions in PDAC relative to those of the coding genome.

## Results

### Clinicopathologic Features of Patient Cohort

The clinical features of this cohort are summarized in Supplementary Table 1. Eight of 10 patients are diagnosed at clinical stage 4 pancreas cancer and two are stage 2B (surgically resectable). Of these latter two patients had a pancreaticoduodenectomy and subsequently developed recurrent metastatic disease. Four patients (PAM01-04) were untreated due to poor performance status at diagnosis and had a short survival (range 0.5-10 months), while six patients (MPAM01-06) had chemotherapy and a relatively longer survival (range 9-49 months). At autopsy, all 10 patients had pathologically confirmed metastases in at least one organ. Nine of ten cases were ductal adenocarcinoma and one case (MPAM02) was undifferentiated carcinoma with osteoclast-like giant cells. Two cases of ductal adenocarcinoma, PAM02 and MPAM06, showed focal squamous differentiation and a third case (PAM01) had neuroendocrine features.

### Mutational Overview of End Stage Pancreatic Ductal Adenocarcinomas

A total of 86 frozen tumor samples and 10 matched normal samples from 10 patients (median 10, range 4-11) were used for this study. We identified a median of 15312 somatic mutations per sample (range 2972-58657) corresponding to median values of 11671 (range 2329 - 37225) single nucleotide variants (SNV), 1903 (213 – 7889) small insertions (INS) and 2664 (430 - 28224) small deletions (DEL) (Supplementary Figure 1) per sample. When these mutations were classified into those occurring in coding, enhancer regions or noncoding-nonenhancer regions (Figure 1a), a median of 119 (range 27 - 434), 91 (16 - 256) and 15075 (2929 - 58216) were identified per region, respectively. We calculated the mutational frequency per million bases (Mb) which revealed a median of 11.9 (range 2.7-43.5), 36.1 (6.4 – 102.1) and 50.7 (9.8 – 196.0) mutations in coding, enhancer and noncoding-nonenhancer regions (Figure 1b). Coding driver gene mutations of *KRAS* (10/10, 100%), *TP53* (7/10, 73%), *CDKN2A* (4/10, 40%), *SMAD4* (4/10, 40%), *ATM*(2/10, 20%), *ARID1A* (2/10, 10%), *ARID2* (1/10, 10%) and *PBRM1* (1/10, 10%) were detected, consistent with findings of previous large cohort studies (Cancer Genome Atlas Research Network. Electronic address and Cancer Genome Atlas Research, 2017)(Nicola Waddell *et al*., 2015) (Figure 1b).

**Fig. 1.**
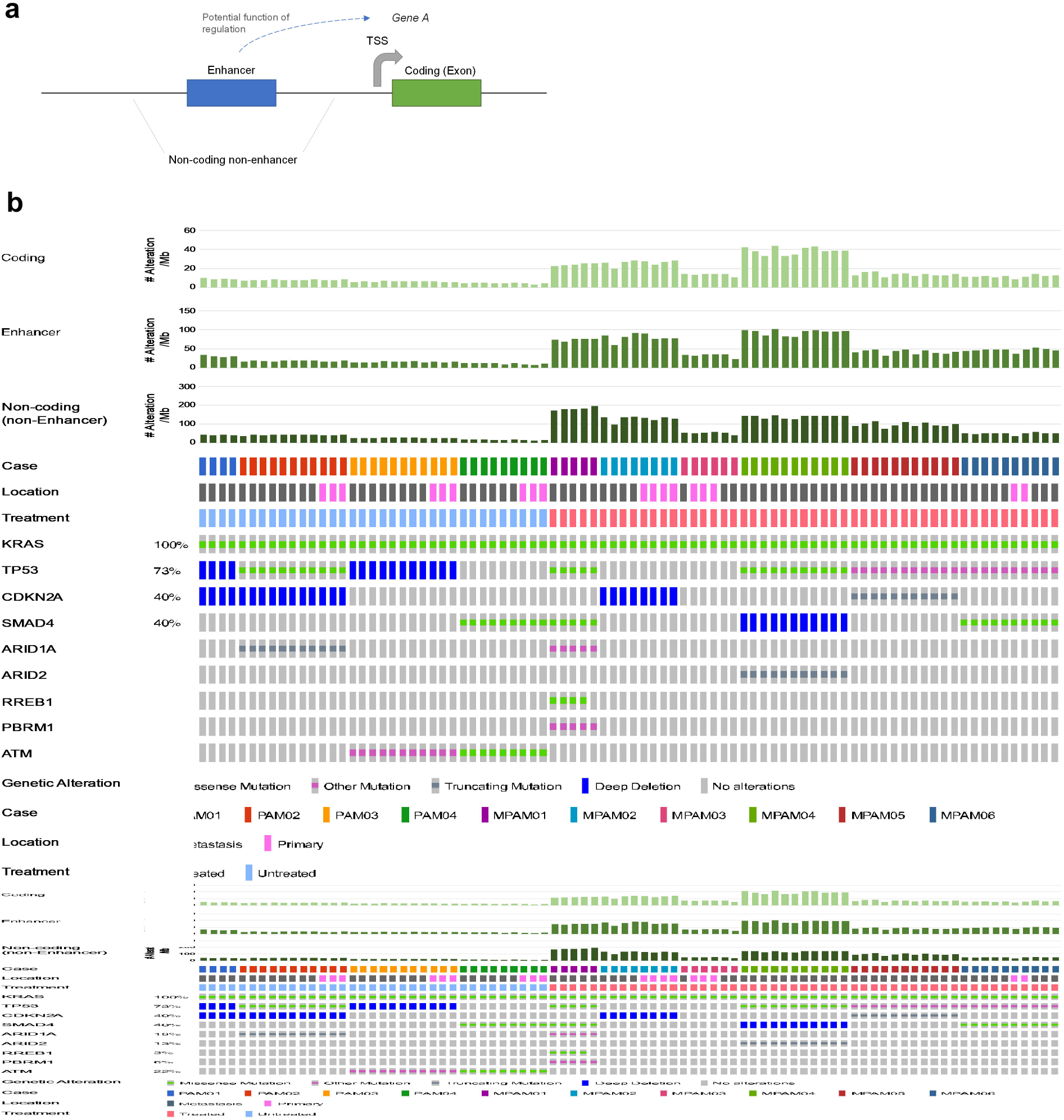
Mutational Overview of End Stage PDACs in Cohort. a. Schematic of coding and noncoding regions studied. b. Oncoprint illustrating the mutational burdens per region of the genome for each sample and the coding driver gene alterations identified per patient. Patients PAM01-PAM04 were treatment naïve and PAM05-PAM10 were treated with chemotherapy with or without radiation.

To understand the relationship between treatment and mutational load we independently evaluated cases with respect to clinical history of receiving chemotherapy. Untreated PDACs had a significantly lower number of mutations (median 8522 mutations per sample, range 2972 – 13266 versus median 32738 per sample, 10968 – 58657) (P < 0.0001, Mann-Whitney U test, two-sided). A similar relationship was found when mutations were categorized into coding, enhancer or noncoding-nonenhancer mutations, as well as when segregated into mutation type (SNV, INS or DEL) (Figure 1b).

### Phylogenic Features of Pancreatic Ductal Adenocarcinomas Studied

Phylogenies were created for each case (Figure 2a, 2b, Supplementary Figure 2, Supplementary Figure 3) and coding driver gene alterations were annotated by LiFD (Reiter *et al*., 2019). The phylogenies of PAM01-PAM04 were previously reported (Makohon-Moore *et al*., 2017) and this current analysis remained entirely consistent with those findings. All common PDAC driver gene alterations (*KRAS, CDKN2A, TP53, SMAD4*) were clonal (truncal) in all ten patients. For six cases multiple regions of the primary tumor were studied in addition to anatomically distinct metastases. In all six cases at least one primary tumor sample was most closely related to a metastasis indicative of subclonal heterogeneity within the primary tumor as previously described (PAM02, PAM03, PAM04, MPAM02, MPAM03, MPAM06).

**Fig. 2.**
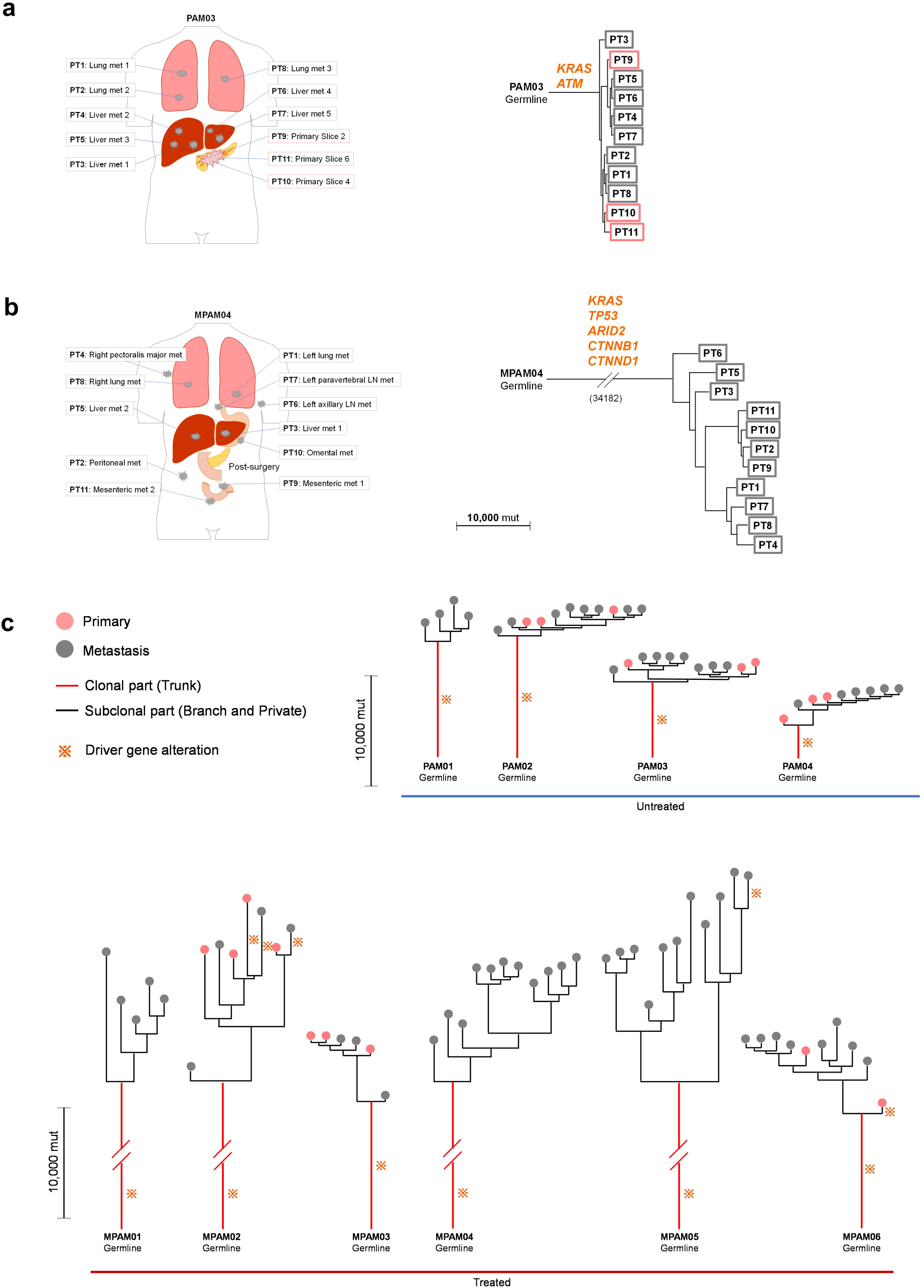
Genetic Evolution of End Stage PDAC. **(a, b)** Anatomical location of autopsy samples and phylogenetic trees in representative untreated (a, PAM03) and treated (b, MPAM04) cases. Driver genes are shown in orange. **(c)** Overview of phylogenetic trees in untreated (PAM01-04) and treated (MPAM01-06) cases. * indicates presence of coding driver gene alteration(s) that are clonal (truncal) or subclonal (branches).

Three PDACs, all treated (MPAM02, MPAM05, MAPM06), had subclonal driver gene mutations in *MUC6, B2M, DNMT3A, or NOTCH2*. Each of these subclonal driver gene alterations was private to a single sample analyzed of the patient, most often the primary tumor (Reiter *et al*., 2019) (Supplementary Table 4), suggestive of treatment induced genetic bottlenecks after metastatic dissemination. Consistent with this interpretation treated PDACs have longer branches compared to untreated PDACs (combination with external and internal branches) (both P < 0.0001, Mann-Whitney U test, two-sided). Collectively, these findings demonstrate this PDAC cohort exhibits the expected clonal dynamics for driver genes and provides the baseline for interpretation of noncoding genetic alterations during subclonal evolution of PDAC (Figure 2c).

### Mutational Characteristics in Pancreatic Ductal Adenocarcinoma

To identify the mutational characteristics of clonal versus subclonal expansions in PDAC, we calculate the relative mutational frequencies in coding, enhancer and noncoding-nonenhancer regions through the entire genome (see methods). In untreated PDACs coding regions had the fewest mutations per MB (median 11.9/Mb), whereas non-enhancer-noncoding regions had the highest number (median 50.7/Mb) (Figure 3a). The difference in relative mutational frequency between coding and enhancer regions was statistically significant for both clonal and subclonal alterations respectively (both P = 0.029, Mann-Whitney U test, two-sided), as were the differences between enhancer and nonenhancer-noncoding regions (P = 0.029 for both clonal and subclonal mutational frequency, Mann-Whitney U test, two-sided). The identical tendencies were observed in treated PDACs (Figure 3b) in that coding regions had fewer mutations per MB than enhancer regions (median 15.7 versus 49.4, P = 0.002, Mann-Whitney U test, two-sided) and enhancer regions had fewer mutations than nonenhancer-noncoding regions (median 49.4 versus 109.3, P = 0.002, Mann-Whitney U test, two-sided). However, pairwise comparisons of coding, enhancer and nonenhancer-noncoding regions between untreated and treated PDACs indicated no differences in mutation frequency. Collectively this suggests two features of the noncoding genome; first, the proportion of mutations in coding, enhancer and non-enhancer regions were preserved despite treatment-induced genetic bottlenecks, and second, enhancers are under negative selection compared to other regions of the noncoding genome.

**Fig. 3.**
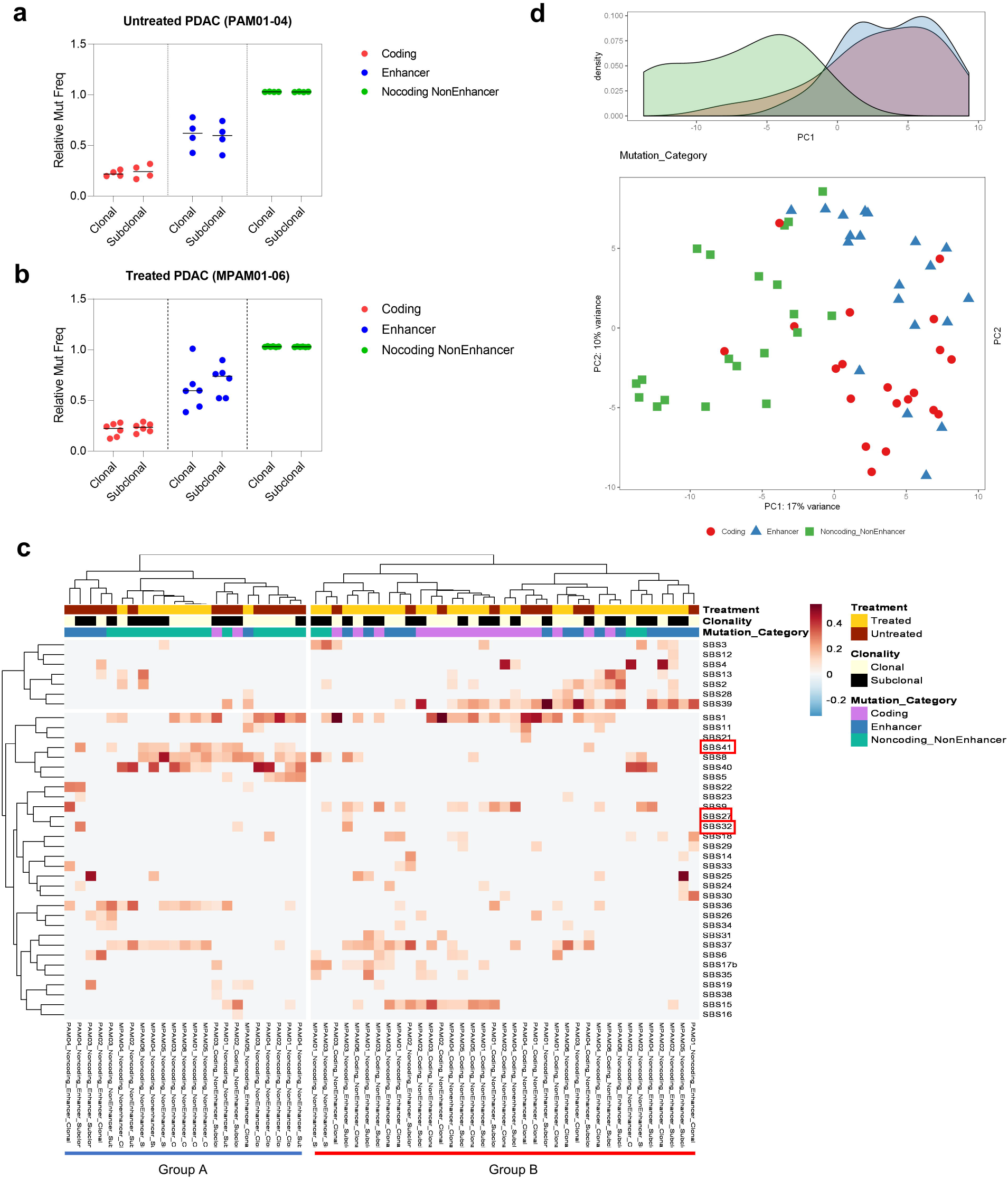
Enhancer Mutation in PDAC. **(a, b)** Relative mutational frequency (mutations/Mb) in coding, enhancer, noncoding-nonenchaner regions in untreated (a) and treated (b) PDAC. **(c)** Heatmap and **(d)** PCA of mutational signature in coding, enhancer, noncoding-nonenhancer regions.

### Mutational Signatures in Pancreatic Ductal Adenocarcinoma

To identify the characteristics of mutational signatures in coding, enhancer and noncoding-nonenhancer regions through the entire genome, we used Palimpsest (Shinde *et al*., 2018) and created a heatmap based on mutational proportions (Figure 3c). Two major clusters were identified based on single base alterations that we designate Group A and Group B based on clustering analysis (see method) for which we annotated with treatment status, clonality and mutation category. Overall, some mutational signatures in coding and enhancer regions were significantly enriched in Group B whereas Group A had more noncoding-nonenhancer mutations associated with a different spectrum of signatures (P < 0.0001, Fisher’s exact test, two-sided). This suggests that enhancers are subject to similar mutational processes as coding DNA than to noncoding-nonenhancer regions. To account for sample size effect (i.e. alteration counts of noncoding-nonenhancer >> coding ≈ enhancer), we performed subsampling analysis using the identical number of mutations (coding counts) for all categories for each case and confirmed the clustering results are similar (data not shown). This tendency was also confirmed in Principal Component Analysis (PCA) based on mutational signatures (Figure 3d) with major overlap found between coding and enhancer alterations in 1^st^ principal component (PC1). Treatment also appeared to affect mutational signatures as samples from untreated patients were relatively increased in Group A whereas treated patients were enriched in Group B (P = 0.020, Fisher’s exact test, two-sided). No grouping differences were found based on clonality (clonal vs subclonal mutations, P = 1.000, Fisher’s exact test, two-sided).

We then focused on the characteristic mutational signatures of each genomic region (coding vs enhancer vs noncoding-nonenhancer), clonality (clonal vs subclonal), or treatment (treated vs untreated). Wilcoxon signed-rank test with FDR (false discovery rate) approach (see method) showed SBS39 was significantly enriched among coding or enhancer alterations (both FDR q-value = 0.017). By contrast, SBS8 (FDR q-value = 0.0007 both in noncoding-nonenhancer vs coding, noncoding-nonenhancer vs enhancer alterations) or SBS40 (FDR q-value = 0.0004 and 0.001 in noncoding-nonenhancer vs coding and in noncoding-nonenhancer vs enhancer alterations) was enriched in noncoding-nonenhancer alterations (Supplementary Table 5). Of note, SBS1, the aged related signature, was found in alterations in coding and noncoding-nonenhancer regions, whereas this signature was an uncommon source of alterations in enhancer regions (FDR q-value = 0.0002 and 0.027 in enhancer vs coding and in noncoding-nonenhancer vs enhancer alterations). These tendencies are confirmed in the International Cancer Genome Consortium (ICGC) cohort (Supplementary Figure 4 and Supplementary Table 6). Overall, we conclude that enhancers are subject to similar mutational processes compared to coding regions.

## Discussion

The evolutionary dynamics of coding DNA in PDAC have been extensively studied. The noncoding genome of PDAC has received less attention, although what has been reported indicates this portion of the genome also has biologic features of significance. For example, Feigin et al has reported recurrent mutations in promoter regions of genes associated with transcriptional regulation (Feigin *et al*., 2017) whereas Rodic et al demonstrated evidence of LINE-1 insertions throughout the somatic genome of primary and metastatic PDAC tissues (Rodić *et al*., 2015). Recently reported pan-cancer studies by the Pan Cancer Analysis of Whole Genomes (PCAWG) Consortium that have included PDAC revealed regions that are significantly affected by recurrent breakpoints, recurrent somatic juxtapositions and recurrent point mutations in non-coding DNA (Rodriguez-Martin *et al*., 2020)(Campbell *et al*., 2020)(Reyna *et al*., 2020)(Rheinbay *et al*., 2020).

We now report several additional features of noncoding DNA in PDAC based on whole genome sequencing. For example, we find that conserved enhancer regions are under negative selection compared to non-coding non-enhancer regions of the genome, with mutational burdens comparable to that of coding DNA. This finding is compatible with another pan-cancer analysis that included PDAC indicating mutations in coding or enhancer DNase I-hypersensitive sites (DHSs) were less than intronic regions (Perera et al., 2016). These differences likely correspond to a number of factors including differences in chromatin structure between enhancers (euchromatin) and intronic DNA (heterochromatin) (Schuster-Böckler and Lehner, 2012) (Polak *et al*., 2015), or differences in DNA repair efficiency among different regions of the genome (Janssen, Colmenares and Karpen, 2018)(Rivera-Mulia and Gilbert, 2016).

MMR can also affect mutational signature in enhancer regions. DNA replication timing is important to determine single-nucleotide substitution patterns in cancer genome (Liu, De and Michor, 2013) and mutational pattern is changed after loss of MMR (Supek and Lehner, 2015). We think MMR susceptibility can cause the differences of mutational signatures between enhancer regions and non-coding non-enhancer regions. In our study, mutational signature in enchanter regions were similar to that in coding regions and different from that in noncoding-nonenhancer regions. Given the fact that MMR prefers exons compared to the introns (Massey and Koren, 2017), this similarity and differences of mutational signature in coding, enhancer and noncoding-nonenhancer regions are understandable. Moreover, based on COSMIC Mutational Signatures (v3 - May 2019) (Alexandrov et al., 2020), Single Base Substitution 39 (SBS39) might a key to understand this difference. This signature was identified more frequently in coding and enhancer mutations (especially found in enhancer mutations) and though the etiology is still unknown, an interesting point was found in the recent PCAWG Consortium (Alexandrov et al., 2020). Though SBS39 was only significant in some types of cancer including pancreas neuroendocrine neoplasm with normal analysis, this SBS had much higher contribution through pan-cancer with additional composite analysis (Alexandrov et al., 2020). We suppose this signature might contribute more than have been thought in previous studies for cancer progression from the point of functional domains in entire genome.

## Methods

### Ethics Statement

This study was approved by the Review Boards of Johns Hopkins School of Medicine and Memorial Sloan Kettering Cancer Center.

### Tissue Samples

A total 86 frozen tumor samples and 10 matched normal samples from 10 patients (median 10, range 4-11) were used for this study. A cohort of four cases from the Gastrointestinal Cancer Rapid Medical Donation Program (GICRMDP) at Johns Hopkins Hospital and six cases from the Last Wish Program (LWP) at Memorial Sloan Kettering Cancer Center were used. Nine patients had a postmortem diagnosis of pancreatic ductal adenocarcinoma and one had a diagnosis of undifferentiated carcinoma with osteoclast like giant cells based on pathologic review. Sections were cut from frozen sections and reviewed to identify those with at least 20% neoplastic cellularity and preserved tissue quality. Normal samples were reviewed to confirm that no contaminating cancer cells were present. Samples meeting these criteria were macro-dissected from serial unstained sections before extraction of genomic DNA using QIAamp DNA Mini Kits (Qiagen).

### Whole Genome Sequencing

Raw data from whole genome sequencing for PAM01-04 were previously generated (Makohon-Moore *et al*., 2017). Genomic DNA was extracted from all tissues for patients MPAM01-06. DNA quantification, library preparation and whole genome sequencing were performed in the MSK Integrated Genomics Operation, and bioinformatics analysis of somatic variants by the MSK Bioinformatics Core. Sequencing, alignment and analysis of MPAM01-06 were performed as described for PAM01-04 (Makohon-Moore *et al*., 2017). Briefly, Illumina HiSeq 2000, HiSeq 2500, HiSeq 4000 or NovaSeq 6000 platform was used to target a coverage of 60X or 80X for tumor and 60X or 30X for normal samples. The resulting sequencing reads were analyzed in silico to assess quality, coverage, as well as alignment to the human reference genome (hg19) using BWA (Li and Durbin, 2009). After read de-duplication, base quality recalibration, and multiple sequence realignment were completed with the Picard Suite and GATK version 3.1 (Depristo *et al*., 2011)(Lisle E. Mose *et al*., 2014), somatic SNVs/INDELs were detected using Mutect version 1.1.6 and HaplotypeCaller version 2.4 (Depristo *et al*., 2011)(L E Mose *et al*., 2014). We excluded low-quality or poorly aligned reads from phylogenetic analysis (Depristo *et al*., 2011)(Cibulskis *et al*., 2013). A median coverage of 76.5x (range 59-124) in tumor and 47.5x (range 28-76) in normal was obtained throughout this cohort.

### Filtering and Annotation of Variants

For all patients, somatic variants were filtered using the following criteria: patient-matched normal coverage ≥10 reads, variant count in patient matched normal ≤2, patient-matched normal variant frequency <0.02, tumor mutant ≥10 reads and tumor variant allele frequency ≥0.05 in at least one tumor sample, under condition with tumor coverage ≥20 reads in all samples). The resulting list of somatic variants were filtered for those present in the coding regions only and subject to further bioinformatic annotation for pathogenicity and germline allele frequencies from healthy populations distributed worldwide using LiFD (Supplementary Table 4) (Reiter *et al*., 2018). Copy number alterations and whole genome duplication were inferred by FACETs (Shen and Seshan, 2016).

### RNA Sequencing

RNA extraction and sequencing performed as recently described (Hayashi *et al*., 2020). Briefly total RNA was extracted using TRIzol (Life Technologies) followed by Rneasy Plus Mini Kit (Qiagen). After library preparation using the TruSeq Stranded Total RNA LT Kit (Illumina catalog # RS-122-1202), samples were barcoded and run on a HiSeq 4000 in a 100bp/100bp or 125/125bp paired end run, using the HiSeq 3000/4000 SBS Kit (Illumina). Output data (FASTQ files) were mapped to the target genome using the rnaStar aligner (version 2.5.0a)(Dobin *et al*., 2013) and postprocessing of the output SAM files was performed using PICARD tools to add read groups and covert it to a compressed BAM format. The expression count matrix from the mapped reads was determined using HTSeq (https://htseq.readthedocs.io/en/release_0.11.1) and the raw count matrix generated by HTSeq was processed using the R/Bioconductor package DESeq2 (http://bioconductor.org/packages/release/bioc/html/DESeq2.html) to normalize the entire dataset between sample groups. Log2 transformated data were used as normalized expression for downstream analyses.

### ICGC Data Set

ICGC pancreas cancer whole genome sequencing data were obtained from ICGC Data Portal (https://dcc.icgc.org/).ICGC pancreatic cancer (version 2016_01_28 for PAAD) RNA-seq data were downloaded through Firebrowse (http://firebrowse.org). Transcripts per million (TPM) was calculated from downloaded RNA-seq data. TPM was used for GSEA and log-2 converted TPM values were used as relative mRNA expression.

### Pancreatic Cancer Enhancer Regions and Enhancer Mutations

Enhancer regions were previously defined based on H3K27ac (activated enhancer histone marks) ChIP-seq data (Johann *et al*., 2016) (Wang *et al*., 2017). Twenty-three H3K27ac ChIP datasets of pancreatic cancer was selected from ChIP atlas (https://chip-atlas.org/) based on number of peaks (> 10000) in combination with manual review on IGV (Supplementary Table 3). All data were merged and analyzed with MACS2 (Zhang *et al*., 2008) and regions with scores over 50 (−10*Log10[MACS2 Q-value]) were identified. These regions were annotated for the closest and second closest genes by GREAT (http://great.stanford.edu/public/html). Regions between 2kb to 50kb from the TSS (transcription start site) were included (all regions farther than 50kb and closer than 2kb from TSS were excluded). Overall 62015 regions were included as “pancreatic cancer enhancer regions” in this study. PDAC enhancer specific regions were identified using Galaxy bedtools ClosestBed (https://usegalaxy.org/) (version 19.09) (Quinlan and Hall, 2010). Enhancer mutations were defined as somatic variants in enhancer regions that overlapped with PDAC enhancer regions using Galaxy bedtools ClosestBed (https://usegalaxy.org/) (version 19.09) (Quinlan and Hall, 2010).

### Evolutionary Analysis

We derived phylogenies for each set of samples by using Treeomics 1.7.9. (Reiter *et al*., 2017). Each phylogeny was rooted at the matched patient’s normal sample and the leaves represented tumor samples. Treeomics employs a Bayesian inference model to account for error-prone sequencing and varying neoplastic cell content to calculate the probability that a specific variant is present or absent. The global optimal tree is based on Mixed Integer Linear programming. All evolutionary analyses were performed based on WGS. Somatic alterations present in all analyzed samples of a PDAC were considered clonal, in a subset of samples or in a single sample considered subclonal.

### Mutational Signature Analysis

Palimpsest (https://github.com/FunGeST/Palimpsest) was used for mutational signature analysis (Shinde *et al*., 2018). All mutations identified in each sample in coding, enhancer, or non-coding non-enhancer regions were used as inputs. Signature proportions in each category of each sample were used for clustering analysis (to divide into two groups; “Group A” and “Group B”) as input to identify characteristics of enhancer region mutational signatures. Mutational signature T-test with FDR approach using t-stage set-up method of Benjamini, Krieger and Yekutieli was performed to identify characteristic signatures between groups.

### Conserved Pancreatic Cancer Enhancer Regions and Gene Annotation

Conserved enhancer regions between humans and mouse were obtained from a previous ENCODE study (Yue *et al*., 2014). Pancreatic cancer enhancer regions with any overlap with ENCODE conserved enhancer regions were defined as “conserved pancreatic cancer enhancer regions”. Out of 62015 pancreatic cancer enhancer regions, 8966 regions (14.5%) passed this criteria. Each conserved enhancer regions and mutations included in these regions (i.e. conserved enhancer mutations) was assigned to the first and second nearest genes within 500kb using GREAT (http://bejerano.stanford.edu/great/public/html/index.php) (version 4.0.4)(McLean et al., 2010).

### Statistics & Reproducibility

All statistics and graphs were performed and made using GraphPad Prism (version 8.2.1) and/or R (version 3.6.1). Parametric distributions were compared by a two-sided Chi Squared test, with correction using a Fisher Exact test for sample sizes <5. Non-parametric distributions were compared using a Mann-Whitney U test (two-sided) and, for analysis of contingency tables, two-sided Fisher’s exact test was used. Each analysis was described in *Results*. Overall survival analyses were performed using the Kaplan-Meier method and curves compared by a log-rank test. Statistical significance was considered if P-value is less than 0.05. FDR q-value was used for GSEA.

No statistical method was used to predetermine sample size. No data were excluded from the analyses as long as the library and/or sequencing quality passed our criteria. The experiments were not randomized. The investigators were not blinded to allocation during experiments and outcome assessment except histological slide review.

## Data and materials availability

DNA sequence data for this study have been deposited at the European Genomephenome Archive (EGA) under accession number EGAD00001006320. The other human resected pancreatic cancer data were derived from the TCGA Research Network: http://cancergenome.nih.gov/. The data-set derived from this resource that supports the findings of this study is available through Firebrowse (http://firebrowse.org/). All data supporting the findings of this study are available upon publication in peer-reviewed journal.

## Acknowledgments

We are grateful to Shinya Oki for his excellent technical support. We gratefully acknowledge the members of the Molecular Diagnostics Service in the Department of Pathology for MSK IMPACT.

## Author contributions

A.H. and C.A.I-D. designed the study; A.H., A.P.M-M., A.Z., M.A.A., P.B., C.A.I-D. collected autopsy samples; A.H. and C.A.I-D. reviewed histology of autopsy samples; A.Z., A.H., A.P.M-M. prepared the DNA samples; A.H., A.P.M-M. and C.A.I-D. performed whole genome sequencing; N.D.S., M.A.A., A.P.M.-M., J.Ho., A.H., J.Hu, and C.A.I-D. analyzed DNA sequencing results and derived the phylogenies; A.P.M-M, J.Ho., A.H., M.P.J. and C.A.I-D. managed the sequencing data; E.M.O. collected clinical information; A.H., and C.A.I-D. wrote the manuscript; all authors reviewed and edited the final manuscript.

## Supplementary Figure Legends

**Supplementary Fig. 1. Mutational Overview of Ten End Stage Pancreatic Cancers.** PAM01-04 are treatment naïve, whereas MPAM01-MPAM06 were treated with chemotherapy.

**Supplementary Fig. 2. Phylogenies of End Stage Untreated PDAC.**

**Supplementary Fig. 3. Phylogenies of End Stage Treated PDAC.**

**Supplementary Fig. 4. Enhancer Mutation in ICGC Cohort PDAC.** Heatmap mutational signature in coding, enhancer, noncoding-nonenhancer regions in ICGC cohort.

