## Supplementary figures and images for "Evolutionary Dynamics of Non-Coding Regions in Pancreatic Ductal Adenocarcinoma"

### Supplementary Figure 1

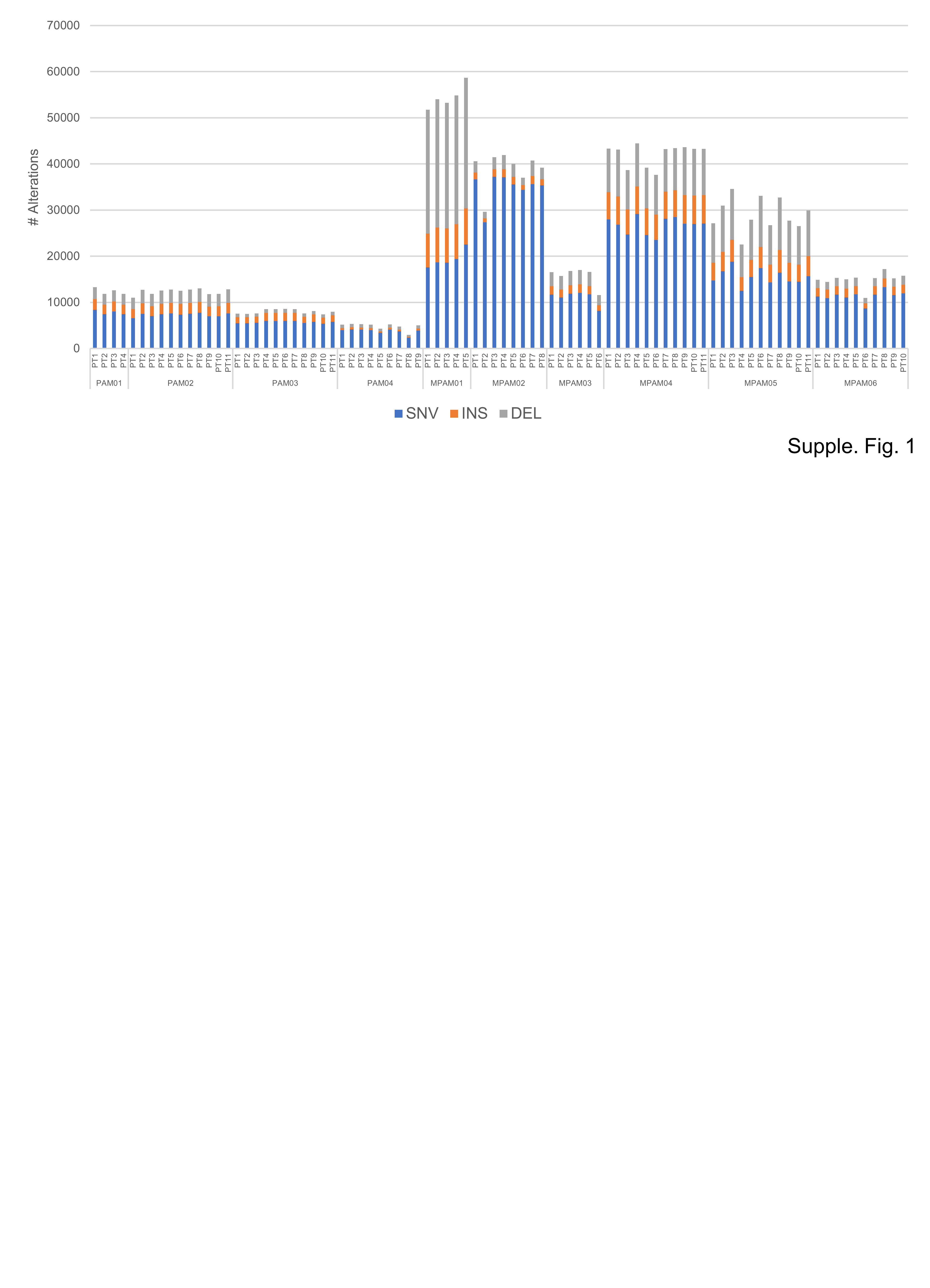

### Supplementary Figure 2

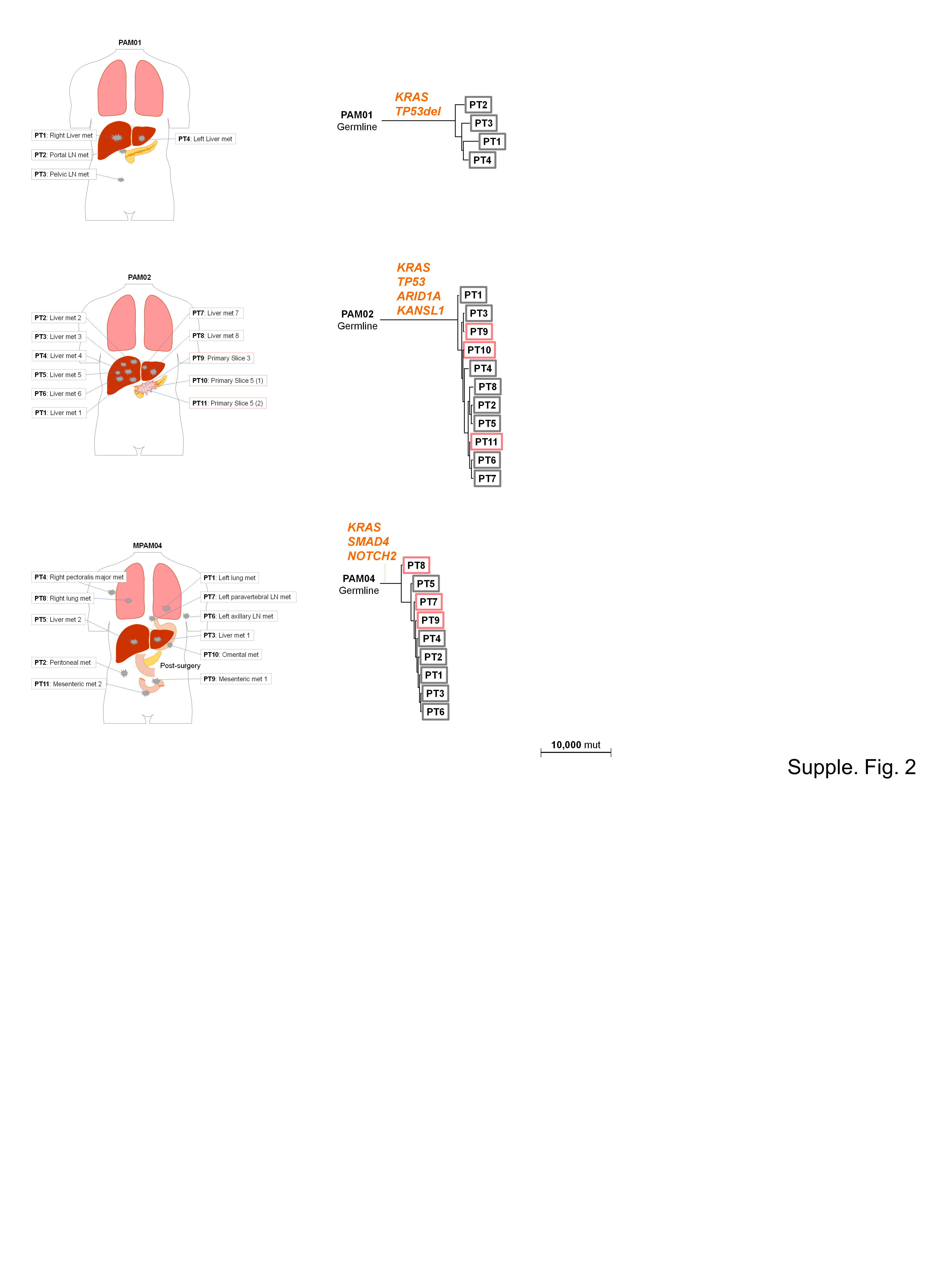

### Supplementary Figure 3

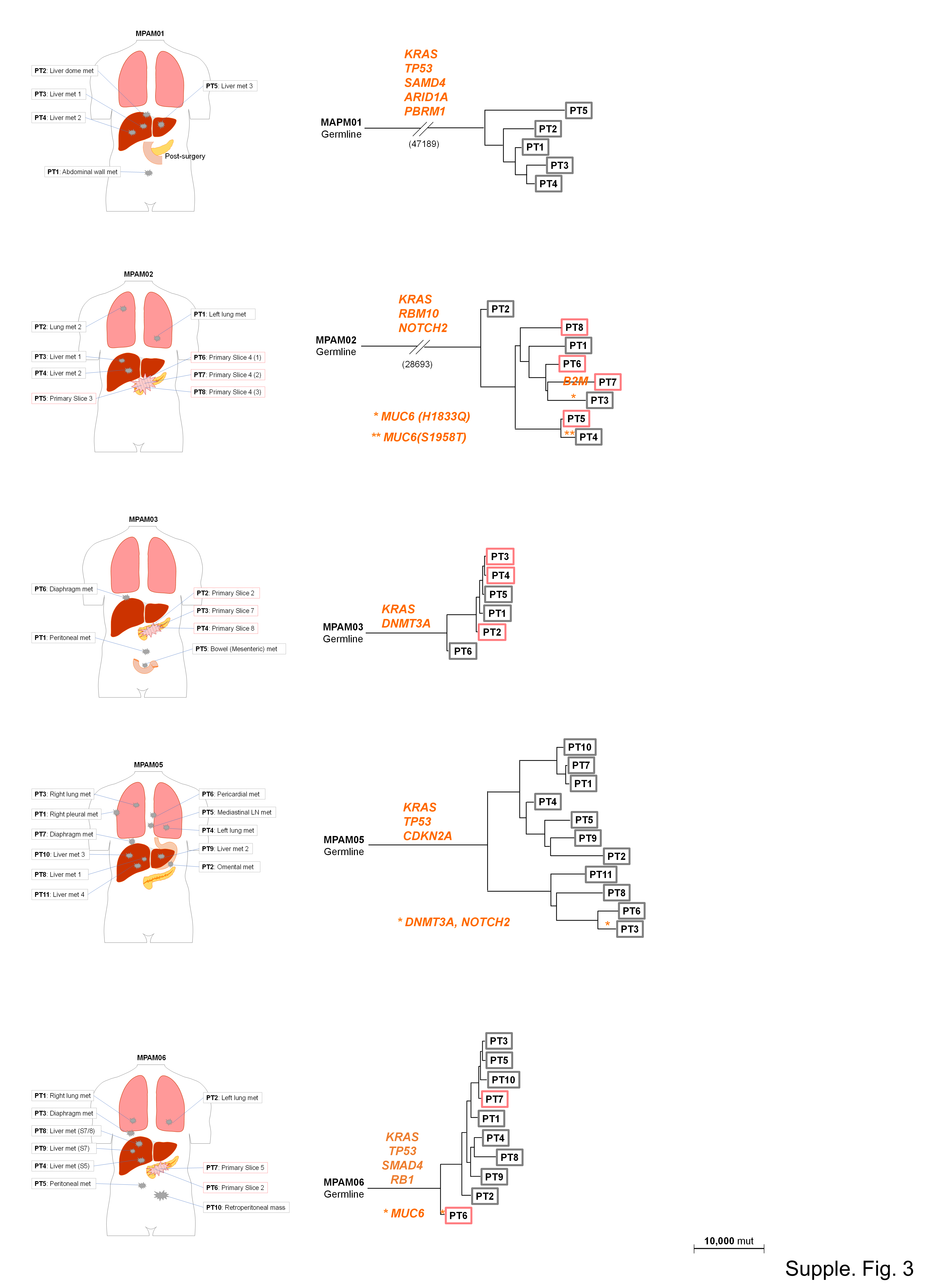

### Supplementary Figure 4

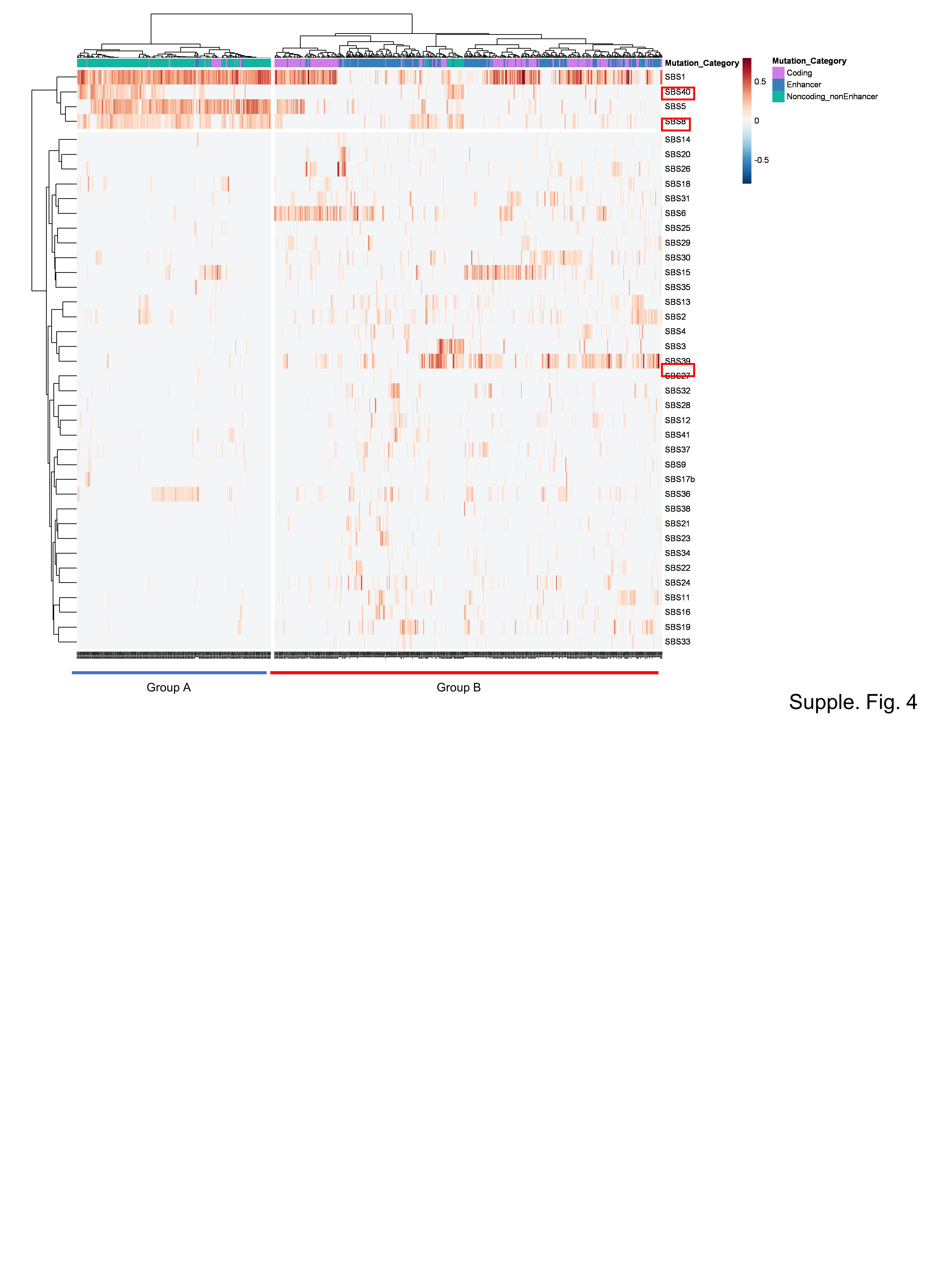
